# Taming a behavioral monster: Resonant song recognition and the evolution of acoustic communication in crickets

**DOI:** 10.1101/2024.07.09.602660

**Authors:** Winston Mann, Bettina Erregger, Ralf Matthias Hennig, Jan Clemens

## Abstract

Rare behavioral phenotypes—behavioral monsters—can challenge hypotheses about the evolution of the neural networks that drive behavior. In crickets, the diversity of song recognition behaviors is thought to be based on the modification of a shared neural network. We here report on a cricket with a novel resonant song recognition pattern that challenges this hypothesis. Females of the species *Anurogryllus muticus* respond to pulse patterns with the period of the male song, but also to song at twice the period. To identify the mechanisms underlying this multi-peaked recognition, we first explored minimal models of resonant behaviors. Though all of the three simple models tested (autocorrelation, rebound, resonate and fire) produced some kind of resonant behavior, only a single-neuron model with an oscillating membrane qualitatively matched the Anurogryllus behavior with regard to both period and duty cycle tuning. Surprisingly, the rebound model, a minimal model of the core mechanism for song recognition in crickets, fails to reproduce the preference for higher duty cycles observed in the behavior, questioning the universality of the core algorithm. However, the behavior is reproduced with a network model that contains all computations from the song recognition network of crickets, revealing the importance of an additional computation not part of the core mechanism: Following the core rebound mechanism in which post-inhibitory rebounds give rise to the resonant period tuning, feed-forward inhibition further shapes the tuning, resulting in the observed behavioral profile. Overall, this shows how unusual behavioral phenotypes can evolve by combining different nonlinear computations at the level of single cells and networks.

## 1 Introduction

Evolution has given rise to diverse animal forms and behaviors. Much of this phenotypic diversity is shaped by the process of mate recognition and sexual selection broadly, and various categories of phenotypic cues—visual, acoustic, chemical, tactile—must be integrated for mate choice decisions to be made. For many species, acoustic signals—calling or courtship songs—are among the first features to be recognized and evaluated in mate choice decisions. The acoustic communication signals produced during courtship behaviors are therefore highly diverse and contribute to species recognition. However, how the neural networks that produce this behavioral diversity evolve is largely unknown. A common hypothesis is that novel behaviors arise from shared neural networks—mother networks—through small changes in connectivity and in cellular properties (Zhu et al., 2023; Bumbarger et al., 2013; Coleman et al., 2023; Ye et al., 2024; Seeholzer et al., 2018). At first sight, the idea of incremental changes in network parameters underlying behavioral evolution is at odds with the observation that behavior can change rapidly (Gallagher et al., 2022; Xu and Shaw, 2021; Ronco et al., 2020; Yona et al., 2018) and outlier species— species with a highly unusual phenotype in a species group—challenge this mother network hypothesis. Evolutionary-developmental biology explains rapid morphological change—so-called “hopeful monsters”—through the re-use and modification of nonlinear gene-regulatory modules (Goldschmidt, 1940; Müller, 2007). Similarly, behavioral innovations—”behavioral monsters”— could emerge from small changes in a network from the nonlinear mapping between the network’s parameters and the behavior.

Experimental tests of the mother network hypothesis are challenging, because they involve characterizing and comparing the network properties across many species in a group and then causally linking the changes in network properties to changes in behavior. However, a precondition for the mother network hypothesis is that the shared network has the capacity to produce the diverse species-specific behaviors in a group. Computational modeling can help assess this capacity from behavioral data, by comparing the observed behavioral diversity with that produced by a computational model of the shared mother network. If the model of the proposed mother network fails to reproduce the behavior of a specific species then that species likely has undergone more drastic changes in its recognition mechanism inconsistent with the mother network hypothesis. Conversely, the hypothesis is supported if the network can reproduce all observed behaviors, including those of the “behavioral monsters”—species with unusual behavioral phenotypes.

We address the question of behavioral diversity and neuronal evolution in the context of acoustic communication in crickets. Males produce pulsed calling songs with species-specific pulse and pause durations ranging between 10 and 80 ms (Fig. 1A, B) (Alexander, 1962). The songs are either produced in chirps consisting of a few pulses or continuously, in trills. Females evaluate the song on the time scale of pulse pause and duration, and of chirps/trills (Fig. 1A) (Grobe et al., 2012). Attractive songs elicit positive phonotaxis in the female. The female tuning for the calling songs can be quantified by measuring the phonotactic behavior for artificial pulse patterns in a two-dimensional parameter space spanned by pulse and pause duration (Fig. 1C). The strength of phonotactic orientation towards the acoustic stimulus then serves as a measure for the strength of recognition. So far, preference functions are known from 18 cricket species, and they all reveal unimodal preferences for a single continuous range of song features (Bailey et al. (2017), Cros and Hedwig (2014), Gray et al. (2016), Hennig (2003), Hennig (2009), Rothbart and Hennig (2012a), Rothbart and Hennig (2012b), Hennig et al. (2016), and Blankers et al. (2015) and Ralf Matthias Hennig, unpublished data). The known preferences fall into three types, characterized by the females’ selectivity for specific features of the pulse song: Tuning for pulse duration, for period (pulse plus pause) and for duty cycle (duration divided by period, referred to from now as “DC”) (Fig. 1F). Tuning for pause duration is a fourth possible phenotype, but this one has not yet been reported in crickets. Song recognition based on the duration or period of acoustic signals is not restricted to crickets but found throughout the animal kingdom (Baker et al., 2019; Araki et al., 2016; Perrodin et al., 2023; Lameira et al., 2024). Understanding the principles underlying the evolution of pulse song recognition in crickets can therefore inform similar studies in others species groups.

**Figure 1:**
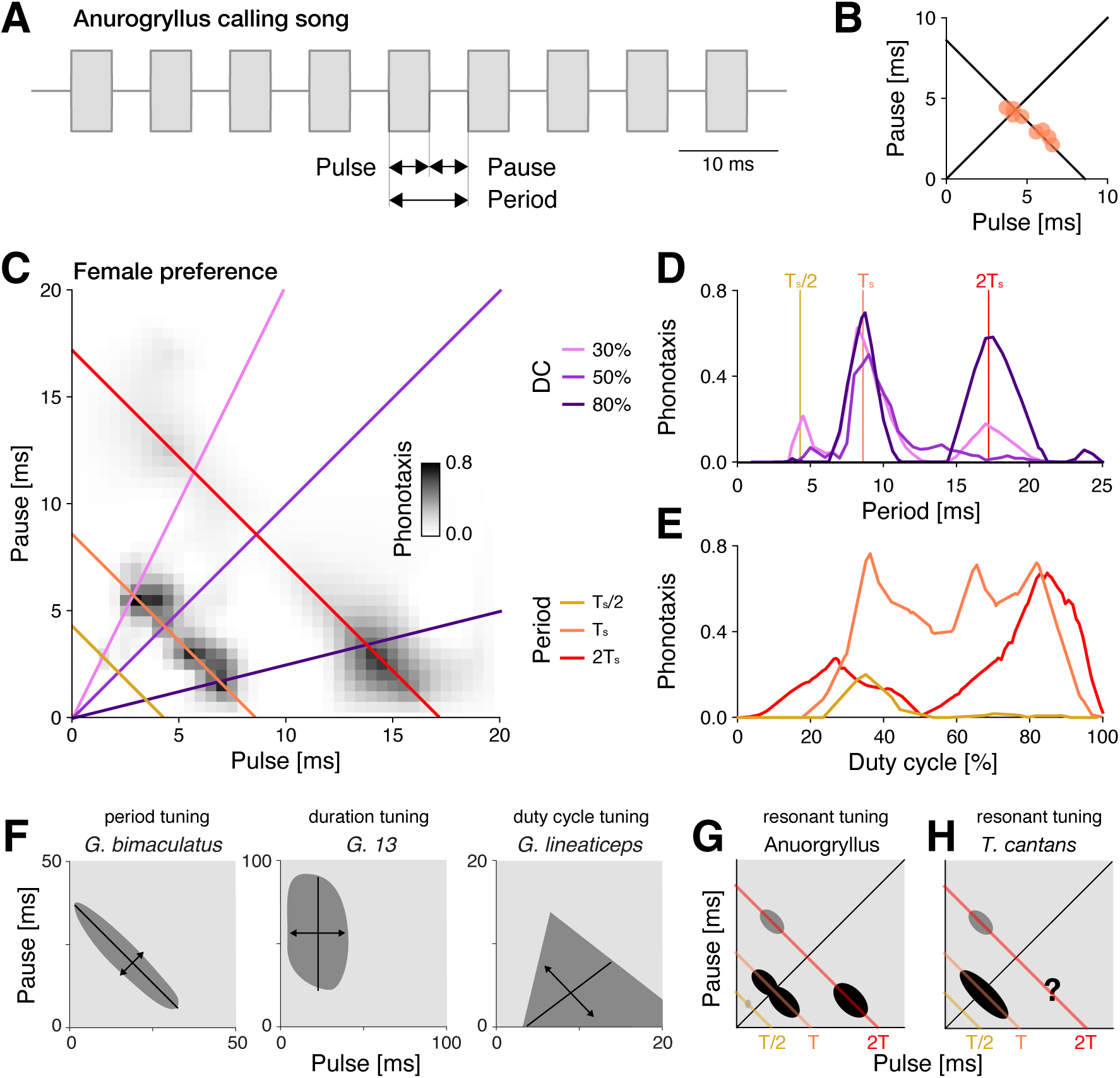
Anurogryllus is a cricket species with resonant song recognition. **A** Schematic of the calling song of males from the cricket species *Anurogryllus muticus* (from now referred to as Anurogryllus). The song consists of a trains of pulses with a specific pulse duration and pause. The period is the sum of pulse and pause and corresponds to the song’s rhythm. The duty cycle (DC) is the percentage of the period occupied by the pulse and corresponds to the song’s energy. **B** Pulse and pause parameters from eight Anurogryllus males. The diagonal line corresponds to a DC of 50%, the antidiagonal to the average pulse period *T_s_* = *8*.*6* ms. See Table 1 for all song parameters. **C** Female phonotaxis for pulse trains with different duration and pause parameters visualized as a pulse-pause field (PPF). Phonotaxis is color coded with darker greys representing stronger phonotactic responses (see color bar). Diagonal lines indicate stimuli with DCs of 30, 50, and 80%, shown in D as the phonotaxis along these diagonals. The anti-diagonal lines show transects with constant period stimuli shown in E at the average pulse period of the male song *T_s_* (orange), at half (*T_s_* /*2*, yellow) and twice (*2 T_s_*, red) the song period. Females respond strongly to pulse patterns with the period of the males’ song, but also at twice that period, indicating resonant song recognition. See table 5 for statistical significance of the individual peaks. The PPF was obtained by interpolation of the average phonotaxis values measured for 75 artificial stimuli in 3–8 females (Fig. S1). **D** Period tuning as a function of DC given by three transects through the PPF in C (see legend in C). Vertical lines indicate *T_s_* /*2* (yellow), *T_s_* (orange), and *2 T_s_* (red). **E** DC tuning as a function of song period, derived from transects through the PPF in C (see legend in C). **F** The three previously known female preference types for the pulse pattern of the male calling song in different cricket species: period (left), duration (middle), and DC (right). The solid black lines indicate the major or most tolerant axis that defines the tuning type, and the double sided arrows perpendicular to the major axis show the most sensitive feature axis. **G** Schematic of the novel resonant recognition from Anurogryllus, simplified from C. **H** Resonant recognition from the katydid *Tettigonia cantans* (Bush and Schul, 2006). The question mark indicates the range of stimulus parameters not tested in the original study. Anti-diagonal lines in G and H indicate stimuli with *T_s_* /*2* (yellow), *T_s_* (orange), and *2 T_s_* (red).

**Table 1:**
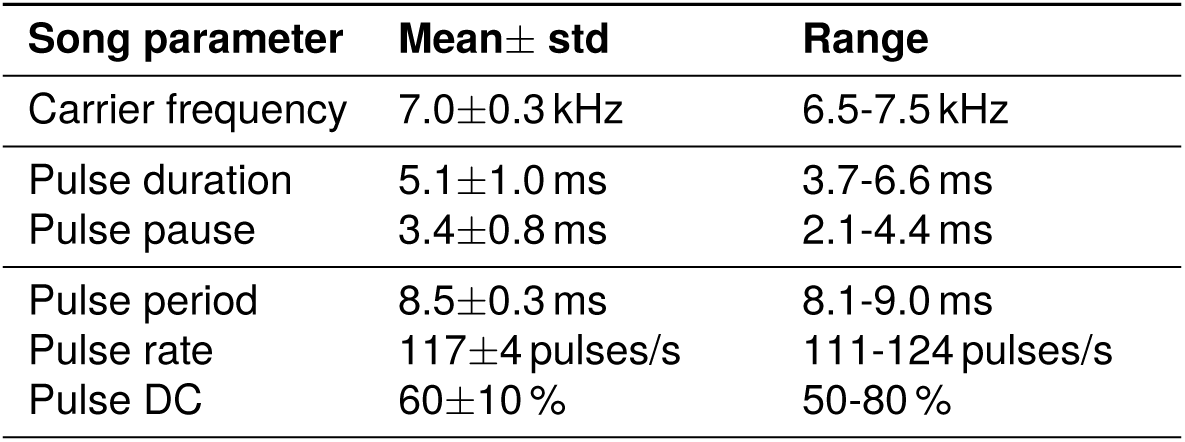
Parameters of the calling song of Anurogryllus males. Data from 8 males over 1 minute of song with at least 7500 pulses per male. Carrier frequencies from (Erregger et al., 2017).

We have recently shown that the song recognition network described in the period-tuned *Gryllus bimaculatus* can produce the diversity of song recognition known in crickets until now. In *G. bimaculatus*, five neurons recognize the song in five steps Schöneich et al., 2015: 1) The ascending neuron 1 (AN1) pools and transmits to the brain information from auditory receptors in the prothorax and produces an intensity-invariant copy of the song pattern (Benda and Hennig, 2008). 2) The local neuron 2 (LN2) receives input from AN1 and provides inhibition to LN5 and LN4. 3) The non-spiking LN5 produces a post-inhibitory rebound potential at the end of each song pulse. 4) LN3 fires only in response to coincident input from the rebound in LN5 and a delayed input from AN1. The input delay from AN1 is tuned such that coincidence only occurs for pulses with the species-specific period of 30 ms. 5) LN4 receives inhibition from LN2, which further sharpens the feature tuning. The tuning of LN4 for pulse song matches that of the phonotaxis response. Similar principles of temporal pattern recognition with delay lines and post-inhibitory rebounds are known from many systems (Carr and Konishi, 1988; Carr, 1993; Kopp-Scheinpflug et al., 2011) and understanding the capacity and constraints of this algorithm in crickets can therefore shed light on temporal pattern recognition across systems. A computational model reproduced the response dynamics of all neurons in this network as well as the behavioral output (Clemens et al., 2020) revealed that the network from *G. bimaculatus* can produce the three preference types known in crickets—preference for period, pulse duration, and duty-cycle—through changes in network parameters like synaptic strengths or intrinsic neuronal properties. Thus, the *G. bimaculatus* network could be the mother network producing the diversity of song recognition in crickets.

We here describe the male song and female preference of a novel cricket species, *Anurogryllus muticus* (from now referred to as Anurogryllus). Anurogryllus is a “behavioral monster” in that it exhibits a multi-peaked recognition phenotype that is novel in crickets and could challenge the hypothesis of a shared mother network: Females are attracted not only to the period of the male song but also to twice the period (Fig. 1C–E). Responses to multiples or fractions of a song’s period have only been shown in a katydid, *Tettigonia cantans* (Fig. 1H), and such responses are consistent with a resonant mechanism for song recognition (Bush and Schul, 2006). Computational modeling in katydids has suggested that delay-based mechanisms can not explain the resonant responses in the katydid and provided evidence for a nonlinear resonant-and-fire (R&F) mechanism of song recognition in katydids (Webb et al., 2007). Importantly, it is unclear whether the computational model of the song recognition network in crickets—which relies on a delay-based mechanism—can produce the resonant preference of Anurogryllus. Thus, Anurogryllus is a challenge to the mother network hypothesis and an opportunity to identify the computational principles that can give rise to resonant tuning.

Here, we provide further support for the mother network hypothesis, by demonstrating that it can produce the resonant recognition behavior of Anurogryllus. We first explore the tuning properties of minimal models of resonant behavior based on network and intracellular mechanisms, and compare these results to those obtained from the full mother network model.

## 2 Results

### 2.1 Anurogryllus demonstrates an unusual resonant recognition phenotype

The calling song of Anurogryllus males consists of continuous trills with a pulse period *T_s_* of *8.5 ± 0.3* ms, which corresponds to a pulse rate *f_s_* of *117.1 ± 4.3* pulses per second (Fig. 1A, B). We refer to the pulse rate measured in pulses per second as f, for notational simplicity. This pulse rate is unusually high for cricket songs, which have pulse rates between 10 and 50 pulses per second (Weissman and Gray, 2019). The song’s DC—given by the ratio of pulse duration and pulse period, and indicating how much of the song is filled by pulses—is 60*±*10% (see Table 1 for a list of all song parameters). To quantify the preference of Anurogryllus females for the calling song we quantified the strength of the females’ phonotaxis response during playback of 75 artificial pulse trains with different pulse and pause duration combinations (Fig. 1C, S1). This confirms that females are attracted (perform positive phonotaxis) to the pulse trains produced by conspecific males: The two-dimensional preference function spanned by pulse duration and pause contains a broad peak covering periods of 8.5 ms and DCs of 33–80%, which overlaps with the distribution of male songs. This peak is partially split along the DC axis (Fig. 1C).

However, the phonotaxis experiments also reveal that females are attracted to songs that differ substantially from the conspecific song and the tuning of these off-target responses implies a resonant recognition phenotype in Anurogryllus (Fig. 1D, E, Table 1). These off-target responses appear at twice or half the song period: First, song with twice the period of the male song (17 ms) with a high DC (90%) is almost as attractive as the conspecific song. Second, females are also weakly attracted to song with twice the conspecific period (17 ms) and lower DC (25%). Lastly, there is a weak and non-significant response peak at half the conspecific period (4.5 ms) and low DC (33%). The responses at integer fractions or multiples of the song’s fundamental rate indicate a resonant response mechanism. If we define *T_s_* = *8.6* ms as the period of the male song, and the fundamental rate *f_s_* = *1* /*T_s_* = *116* pulses per second, then the weak peak at half the period, *T_s_* /*2 ≈ 4.3* ms, corresponds to the second harmonic, *2 f_s_*, while the peaks at twice the period, *2 T_s_ ≈ 17.2* ms, correspond to the second subharmonic, *f_s_* /*2*.

So far, a resonant song recognition behavior—with responses to three different types of pulse patterns—has not been reported in a cricket (Fig. 1F–G)—it was previously known only in the katydid *Tettigonia cantans* (Bush and Schul, 2006) (Fig. 1H). The resonant phenotype in *T. cantans* is similar to that of Anurogryllus: *T. cantans* females are attracted to pulse trains with the period of the male song (period 40 ms, DC 50%), and to subharmonics of the male song—songs with twice the conspecific period (80 ms, DC 25%). *T. cantans* does not respond to harmonics (half the period, 20ms) and it was not tested whether females are attracted at twice the period at higher DCs, the pattern that Anurogryllus is most responsive to apart from the conspecific song. A simple delay-line based mechanism in *T. cantans* was ruled out as a potential mechanism for resonance using experimental tests, but a resonate-and-fire neuron model with oscillatory membrane properties could reproduce the resonant song preference (Bush and Schul, 2006; Webb et al., 2007). Oscillatory neurons have therefore been proposed as a mechanism for song recognition in *T. cantans*. However, the rebound-based mechanism at the core of the song recognition network in crickets had not been considered, and it is unclear whether oscillatory neurons can reproduce the particular pattern of resonance observed in Anurogryllus.

### 2.2 Simple models provide insight into computational mechanism of resonant tuning

The resonant phenotype in Anurogryllus challenges the mother network hypothesis, as the model of the song recognition network in crickets was only shown to produce all known single-peaked phenotypes, not the specific resonant phenotype of Anurogryllys (Clemens et al., 2021) (Figs 1F, G). We therefore tested whether this model network could also produce the resonant tuning of Anurogryllus. However, given that the computational model of the song recognition network in crickets is complex and has many parameters, we decided to first identify the computational principles and constraints that shape resonant tuning by investigating the ability of simple network and single-neuron models to qualitatively reproduce the resonant behavior of Anurogryllus. Simple models allow us to 1) isolate the minimal set of computations required for generating resonant behaviors, 2) facilitate the interpretation of the more complex network model, and 3) rule in or out alternative mechanisms not currently part of the mother network but that might be easily acquired during evolution. Given the simplicity of the models chosen, our goal was not a detailed reproduction of the Anurogryllus behavior (Fig. 1C), but a reproduction of the most prominent properties of the period and DC tuning: namely the broad DC peak at the period of the male song, *T_s_*, and the two response peaks at *2 Tf_s_*, with the dominant peak at high DC (Fig. 1G).

We fitted three simple models to the behavioral data from Anurogryllus: First, an autocorrelation model, which consists of a delay line and a coincidence detector (Bush and Schul, 2006). This is the simplest model that can produce resonances and shows that delays alone can produce resonant response peaks. Second, the rebound model, which is an extension of the autocorrelation model and captures the core computation of the mother network, in which the non-delayed input to the coincidence detector consists of offset responses from a post-inhibitory rebound (Schöneich et al., 2015; Clemens et al., 2021). The rebound model will reveal whether the core-computation of the mother network—a delay line, rebound, and coincidence detection—is sufficient to produce the resonant tuning of Anurogryllus. Lastly, we examined the a resonate-and-fire (R&F) neuron, a single-neuron model with subthreshold membrane oscillations that reproduced the resonant behavior of *T. cantans* (Izhikevich, 2001; Webb et al., 2007). This last model will allow us to examine how changes in intracellular properties, rather than network properties, can produce the resonant song recognition of Anurogryllus.

#### Autocorrelation models produce resonant tuning but do not match the Anurogryllus behavior

In an autocorrelation model, the song input is split into two pathways, one with a delay *Δ_ac_*, and one without a delay (Fig. 2A). Responses from the delayed and non-delayed pathways are then multiplied in a coincidence detector, that only responds when the delayed and the non-delayed inputs overlap in time. The model response is then taken as being proportional to the average output of the coincidence detector over the song.

**Figure 2:**
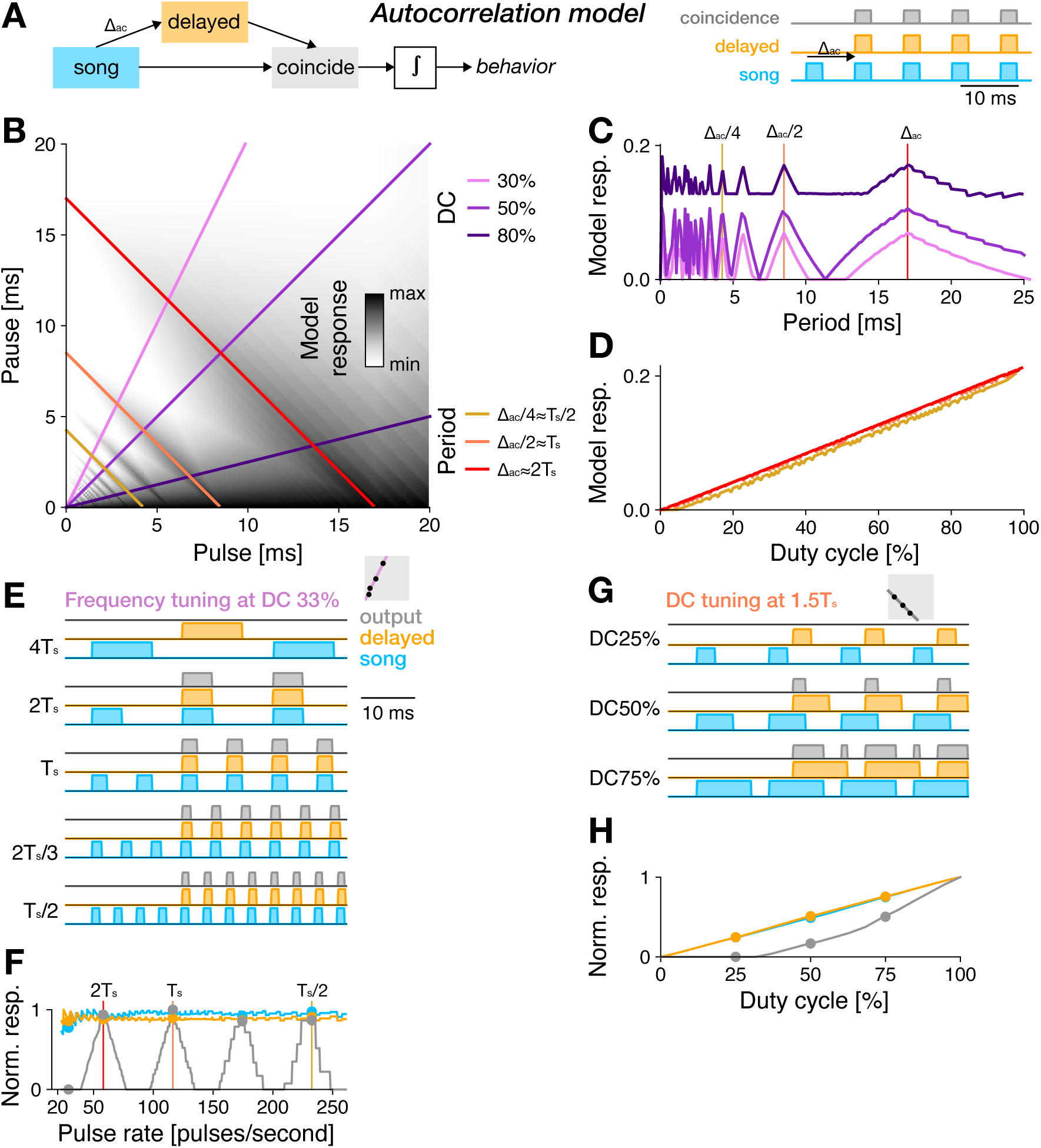
An autocorrelation model produces resonant tuning. **A** In the autocorrelation (AC) model, a non-delayed (blue) and delayed (orange) copy of the stimulus are multiplied in a coincidence detector (grey). The output of the coincidence detector is integrated over the stimulus to predict the model response. The example traces show coincidence for song with a pulse period that equals the delay *Δ_ac_*. **B** PPF for the autocorrelation model fitted to the preference data in 1C. Predicted response values are coded in greyscale (see color bar). Colored lines correspond to the DC and period transects shown in C and D (see legend). **C** Period tuning of the autocorrelation model for different DCs (see legend in B). Resonant peaks arise at even and odd fractions of the delay parameter *Δ_ac_ ≈ 2 T_s_*. Vertical lines indicate the pulse periods transects shown in B. **D** DC tuning for three different pulse periods (see legend in B), corresponding to *T_s_* /*2*, *T_2_*, and *2 T*. DC tuning is high-pass for all periods. **E** Response traces from the autocorrelation model for songs with different periods (fractions and multiples of *T_s_*) and a DC of 33%. Resonant peaks arise from coincidence at integer fractions (e.g. *1 Δ_ac_* /*2* = *2 T_s_* /*3*) but not at multiples (*2 Δ_ac_* = *2 T_s_*) of the delay parameter (stimulus–blue, delayed stimulus–orange, response–grey, see legend to the right). **F** Pulse rate tuning given by the integral of the stimulus (blue), the delayed stimulus (orange), and the response (grey) at 50% DC. Response peaks arise at integer multiples fractions of *Δ_ac_*. Dots indicate pulse patterns shown in E. Vertical lines indicate the song periods shown in D. **G, H** Response traces for different DCs (25, 50, 75%) (G) and DC tuning (H) at a non-resonant pulse rate (*1.5 T_s_* = *12.9* ms). Increasing the DC leads to coincidence even at this non-resonant pulse rate. Same color code as in E, F. Gray boxes in E and G illustrate the stimulus parameters for which traces are shown in the context of the PPF (compare B).

The autocorrelation model fitted to the Anurogryllus data produces resonant response peaks for pulse rates at integer fractions, but not at multiples, of the delay *Δ_ac_* (Fig. 2B, C). The fitted value of *Δ_ac_* = *17* ms corresponds to *2 T_s_*, the peak at twice the pulse period in the behavioral data (Fig. 1E). Coincidence occurs if *nT* = *Δ_ac_*, leading to resonant peaks at periods that are fractions of the delay *T* = *Δ_ac_* /*n* (or at pulse rates *f* = *n*/*Δ_ac_*) (Fig. 2E, F). Thus, resonant peaks in the autocorrelation model arise at even and odd fractions of *Δ_ac_* and coincide with *T_s_* and *2 T_s_*. However, the behavior only exhibits responses at even fractions of *Δ_ac_*. The lack of peaks at odd fractions of *Δ_ac_* in Anurogryllus renders a pure autocorrelation-based mechanism for song recognition unlikely (Fig. 1E).

Similar to the period tuning, the DC tuning of the fitted autocorrelation also does not match the behavioral data: The output of the autocorrelation model increases linearly with DC (Fig. 2D), with maximal responses for constant tones without a pause (DC 100%). By contrast, Anurogrullus exhibits complex DC tuning with multiple peaks and, importantly, does not respond well to pulse trains with very high DCs (Fig. 1E). The DC bias in the autocorrelation model arises because songs with longer pulses and shorter pauses are more likely to produce coincidence for any given delay (Fig. 2F, Fig. 2G, H).

In sum, the autocorrelation model demonstrates that a delay is sufficient to produce resonance. However, autocorrelation alone is insufficient to qualitatively reproduce the pulse rate and DC tuning found in Anurogryllus.

#### A rebound mechanism suppresses responses to pulse trains with high duty cycles

The core computation for song recognition in the cricket *G. bimaculatus* is an extension of the autocorrelation model (Schöneich et al., 2015; Clemens et al., 2021) (Fig. 3A): As in the autocorrelation model, the song is split into a delayed and a non-delayed path. The non-delayed path is then sign-inverted and filtered to produce transient responses at the end of each pulse, to mimic a post-inhibitory rebound. The rebound model produces outputs only when the delayed input coincides with the rebound.

**Figure 3:**
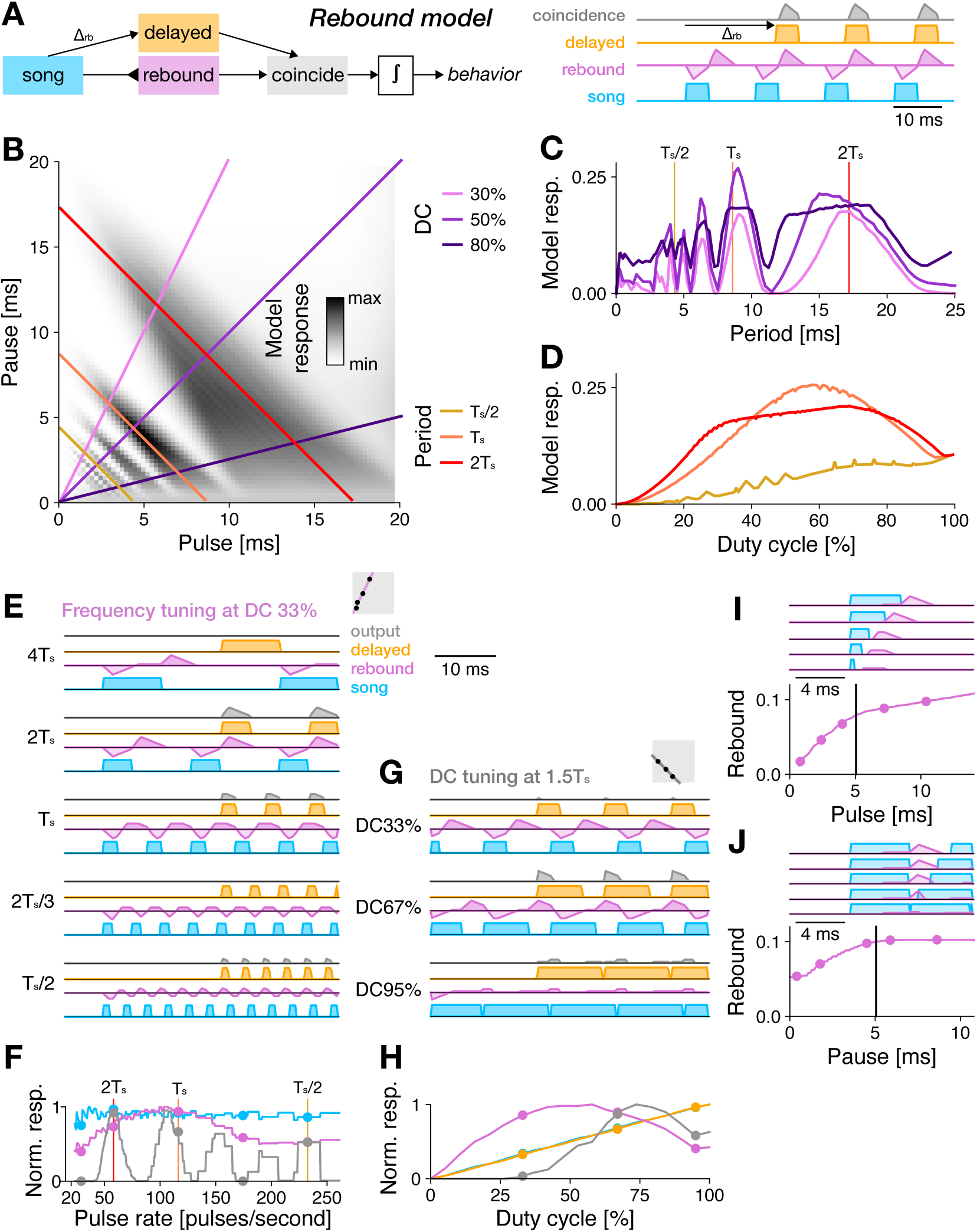
Tuning for pulse rate and duty cycle in the rebound model fitted to Anurorgryllus behavior. **A** The rebound model is an extension of the autocorrelation model. The non-delayed branch (purple) is sign-inverted (blunt ended arrow indicates inhibition) and filtered by a bi-phasic filter to produce transient responses at pulse offsets that mimic a post-inhibitory rebound. The positive part of the rebound and the delayed stimulus are then combined through coincidence detection. **B** PPF for the rebound model fitted to the preference data in 1C. Predicted response values are color coded (see color bar). Colored lines correspond to the DC and period transects shown in C and D (see legend). **C** Period tuning of the rebound model for different DCs (see legend in C). Vertical lines correspond to the pulse period transects shown in B. **D** DC tuning for three different pulse periods (see legend in C). DC tuning is high-pass for short periods (*T_s_* /*2*, yellow) and band-pass for intermediate and long periods (*T_s_* (orange), *2 T_s_* (red)). **E** Response traces of the rebound model for songs with different periods (fractions and multiples of *T_s_*) and a DC of 33% (stimulus–blue, rebound response–pink, delayed stimulus–orange, response–grey, see legend to the right). **F** Pulse rate tuning given by the integral of the stimulus. Response peaks arise at integer multiples of f. Dots indicate periods shown in E. **G, H** Response traces for a DC sweep (33, 67, 95%) (G) and DC tuning (H) at a non-resonant period of *1.5 T_s_* = *12.9* ms. Even at this non-resonant period, responses increase with DC, consistent with the broadening of the response peaks with DC in B and C. Responses decrease at very high DCs (short pauses), because the rebound is truncated by the next pulse (see J). Same color scheme as in E, F. **I, J** Integral of the rebound as a function of pulse duration (I) and pause (J). Dots in the curves (bottom) indicate example traces shown on top of each curve. A minimum pulse duration and pause duration (black lines) are required for the rebound to fully develop. At short pauses the rebound is interrupted by the following pulse (J). Gray boxes in E and G illustrate the stimulus parameters for which traces are shown in the context of the PPF (compare B).

The pulse rate tuning of the rebound model resembles that of the autocorrelation model, with resonant peaks arising close to even and odd fractions of the delay *Δ_rb_* (Fig. 3B, C, compare 2C). However, the fitted value of *Δ_rb_* = *23 ms* matches neither multiples nor fractions of *T_s_*. This is because the rebound is produced at the end of each pulse and coincidence therefore occurs if *n · T* + *D* = *Δ_rb_*, where *D* is the pulse duration (Fig. 3E, F). Resonant peaks occur at *T* = (*Δ_rb_ − D*)/*n* (equation 2.2) or *f* = *n*/(*Δ_rb_ − D*), close to even and odd fractions of *2 T_s_* (Fig. 3A, B, E, F). The responses to odd fractions of *T_s_* in the rebound model are not found in the behavioral data. Therefore, a pure rebound mechanism is unlikely to produce the Anurogryllus behavior.

The DC tuning of the rebound model is band-pass, with reduced but non-zero responses for continuous tones (high DC) (Fig. 3D, G). This band-pass tuning arises from two opposing processes: On the one hand, responses increase with pulse duration up to a point set by the duration of the inhibitory filter lobe that produces the rebound. This is because the rebound is strongest if the pulse is long enough to saturate the rebound, which happens when it fully overlaps the inhibitory filter lobe (Fig. 3I). However, a further increase in pulse duration at a fixed pulse period shortens the pauses and for short pauses, the rebound is interrupted by the next pulse (Fig. 3G, J).

Overall, the rebound model fails to reproduce the qualitative features of the Anurogryllus responses. Period tuning exhibits excess peaks at odd fractions of the pulse rate as in the autocorrelation model. While the DC tuning is band-pass, as in Anurogryllus, responses to constant tones are still evident and the characteristic pattern with split-peaks is missing. This failure to reproduce the Anurogryllus behavior is surprising given that the rebound constitutes the core mechanism of song recognition in crickets. However, we will show below that a rebound mechanism can produce the Anurogryllus behavior when combined with other computations found in the full network.

#### The resonate and fire model is a simple model that qualitatively matches the Anurogryllus behavior best

As the last simple model, we fitted a resonate-and-fire (R&F) model to the Anurogryllus data. In contrast to the autocorrelation and rebound models, which are network models, the R&F model is a single neuron model that consists of coupled current and voltage-like variables (Fig. 4A) (Izhikevich, 2001). This coupling leads to input-driven damped oscillations with a characteristic frequency *f_r_*_&*f*_. Inputs that arrive at positive/negative phases of the oscillation amplify/suppress this oscillation. If the voltage reaches a threshold, a spike is elicited and the current and voltage are reset. The R&F model can produce resonant behavior if the damping is weak and it was used to reproduce the resonant song recognition from *T. cantans* (Bush and Schul, 2006; Webb et al., 2007).

**Figure 4:**
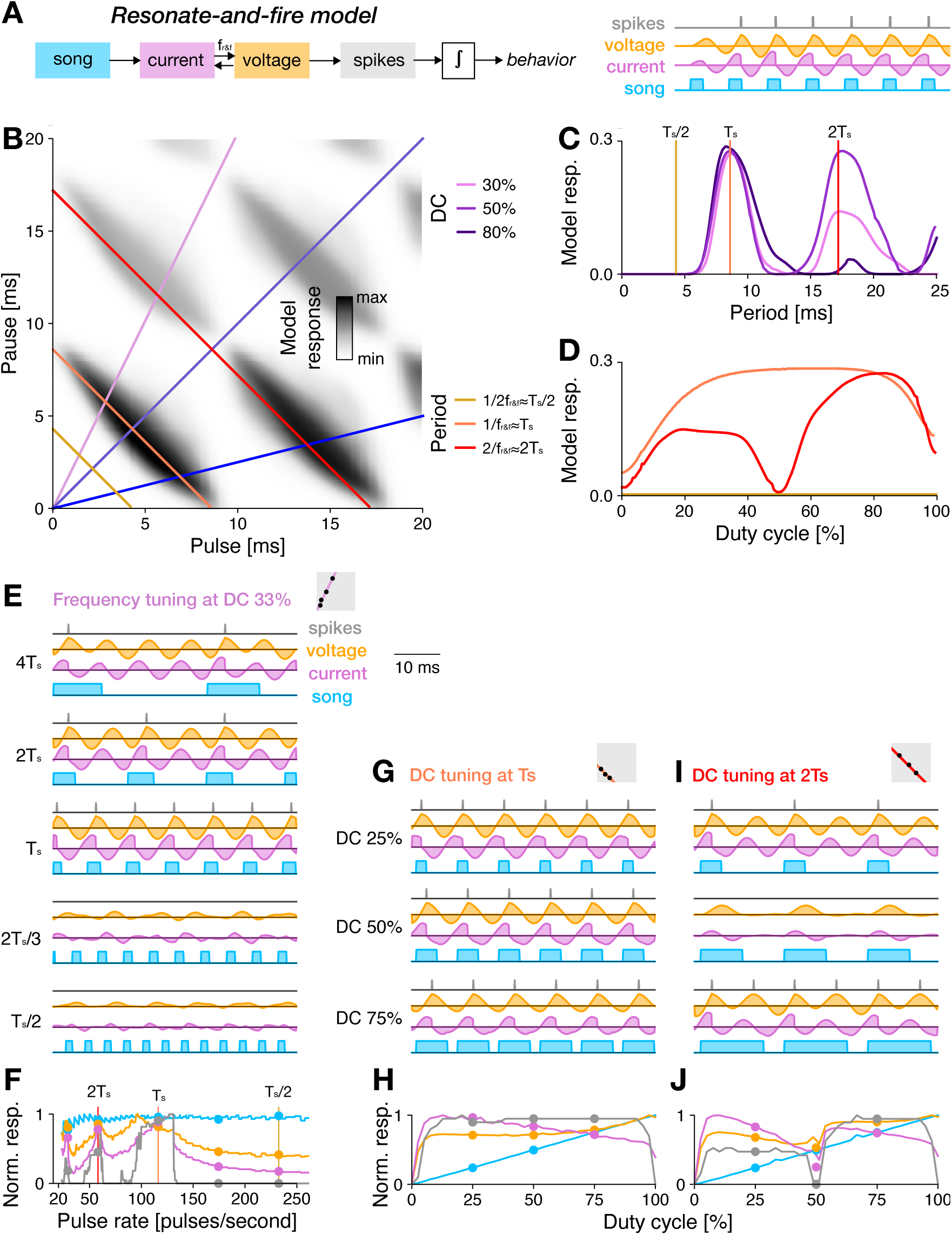
Tuning for pulse rate and duty cycle in the resonate and fire model fitted to Anurogryllus behavior. **A** The resonate-and-fire (R&F) model is a spiking neuron model with bidirectionally coupled current (purple) and voltagelike (orange) variables. Inputs currents trigger oscillations with a frequency *ω*. Inputs are excitatory during positive phases and inhibitory during negative phases of the oscillations. If the voltage exceeds a threshold, a spike (grey) is elicited and the current and voltage are reset. **B** Pulse-pause field (PPF) for the R&F model fitted to Anurogryllus data. Colored lines correspond to the DC and period transects shown in C and D (see legend). **C** Period tuning of the R&F model for different DCs. Resonant peaks arise at periods at integer multiples of *T_s_*. The response at *2 T_s_* is attenuated for lower DCs, as in the behavior. Vertical lines correspond to the frequencies shown in D. **D** DC tuning for three different pulse periods. There is no peak for *T_s_* /*2*. At *T_s_*, the DC tuning is band-pass. At *2 T_s_*, the DC tuning is bimodal, as in the data. **E** Response traces for the R&F model for songs with different periods (fractions and multiples of *T_s_*) and a DC of 33% (stimulus– blue, current–pink, voltage–orange, spikes–grey, see legend). Membrane oscillations and responses are weak at fractions at *T_s_*. Responses are strong at integer multiples of *T_s_*. **F** Pulse rate tuning at DC 33%. Shown are the integrals of the stimulus (blue) and spiking response (grey). The current-like (pink) and voltage-like (orange) variables were rectified before integration. **G, I** Response traces for different DCs at *T_s_* (G) and *2 T_s_* (I). **H, J** DC tuning at *T_s_* (H) and *2 T_s_* (J). Dots mark the stimuli shown in G and I. DC tuning is unimodal at *T_s_* and bimodal at *2 T_s_*. Gray boxes in E, G, and I illustrate the stimulus parameters for which traces are shown in the context of the PPF (compare B).

The R&F model fitted to Anurogryllus data is weakly damped (less than 2% of the stimulus gain). It has a characteristic frequency *f_r_*_&*f*_ = *109* Hz, which translates to *T_r_*_&*f*_ = *9.2* ms—close to the pulse period of the Anurogryllus song. The R&F responds strongly if the incoming pulses hit the intrinsic oscillation during excitatory phases and resonant peaks therefore arise at multiples of the period *n · T_r_*_&*f*_ = *n*/*f_r_*_&*f*_. Thus, contrary to the autocorrelation and rebound models, the R&F model responds only to multiples (subharmonics), but not to fractions of *T_r_*_&*f*_ (harmonics) (Fig. 4B, C). In the R&F, responses to fractions of *T_r_*_&*f*_ are suppressed, because inputs faster than *T_r_*_&*f*_ will arrive not only during the excitatory but also during the inhibitory phase of the intrinsic oscillation, reducing the net drive to the spike generator. By contrast, responses at multiples of *T_s_* exist because subsequent pulses will arrive during the excitatory phase of the membrane oscillation (Fig. 4E, F). The R&F model produces the Anurogryllus responses at *T_s_* and *2 T_s_* and, apart from excess responses at higher multiples of *T_s_*, matches the behavior well.

The DC tuning of the R&F model is more complex than that of the autocorrelation and rebound models. At *T_r_*_&*f*_, the model responds with a single, broad peak to different DCs, whereas at *2 T_r_*_&*f*_, two separate peaks—at high and low DCs—are visible (Fig. 4B, D). The peak at the higher DC is greater than that at the lower DC, consistent with the Anurogryllus behavior. With this DC tuning, the R&F model qualitatively matches the Anurogryllus data best out of all models tested so far (Fig. 1F). Note that the R&F also produces peaks at *3 T_r_*_&*f*_, but stimuli covering these periods were not tested experimentally.

How does this complex DC tuning arise in the relatively simple R&F model? In the model, inputs during the excitatory phase of the membrane potential oscillation amplify the oscillation and therefore elicit spiking responses, while inputs during the negative phase suppress the spiking responses. Songs with a pulse period of *T_s_* match the period of the membrane oscillation and an input with a DC of 50% will produce the maximum output because it covers only the excitatory phase of the oscillations (Fig. 1G, H). Shorter pulses (DC<50%) will produce weaker voltage responses because they engage the excitatory phase less, and longer pulses (DC>50%) will produce weaker voltage responses because they extend into the inhibitory phase. Pulse patterns with a pulse period of *2 T_s_* —twice the period of the oscillation—produce DC tuning with two broad peaks—around DC 25% and around DC 75%—and no responses at DC 50% (Fig. 1I, J). The responses at DC 50% are suppressed because the pulse covers one full period of the oscillation, and therefore equally engages the excitatory and the inhibitory phases of the oscillation, resulting in weak spiking responses. Stimuli with smaller or larger DCs produce stronger responses because more of the excitatory phases of the oscillation are engaged. The peak at higher DCs is higher than that at lower DCs because the pulse hits the excitatory phase once per period for DCs below 50% and twice for DCs above 50% (Webb et al., 2007), as in (Fig. 4I).

#### Simple network models, unlike the single neuron model, fail to reproduce the behavioral period and DC characteristics

Overall, none of the simple models were able to fully reproduce the Anurogryllus tuning. However, a single-neuron model—the R&F model—came closest, suggesting that changes in single neuron properties might underlie the emergence of resonant tuning in Anurogryllus (Fig. 4). By contrast, simple delay-based models (autocorrelation and rebound) are insufficient to recover the Anurogryllus tuning (Figs 2, 3): The delay-based models are resonant but they produce strong responses to very short periods (fractions of *T_s_*) and are unable to replicate the DC tuning of Anurogryllus, in particular the double-peaked DC tuning at *2 T_s_*. Importantly, the failure of the rebound model, which replicates the hypothesized core mechanism of song recognition in crickets, challenges the mother network hypothesis (Schöneich et al., 2015; Clemens et al., 2021)). However, the mother network, developed using electrophysiological data from *G. bimaculatus*, contains additional computations like adaptation and feed-forward inhibition. We therefore fitted a model of the full network, previously developed in Clemens et al. (2021) (Fig. 5A), to the behavioral data from Anurogryllus to test whether these additional computations can produce the behavior.

**Figure 5:**
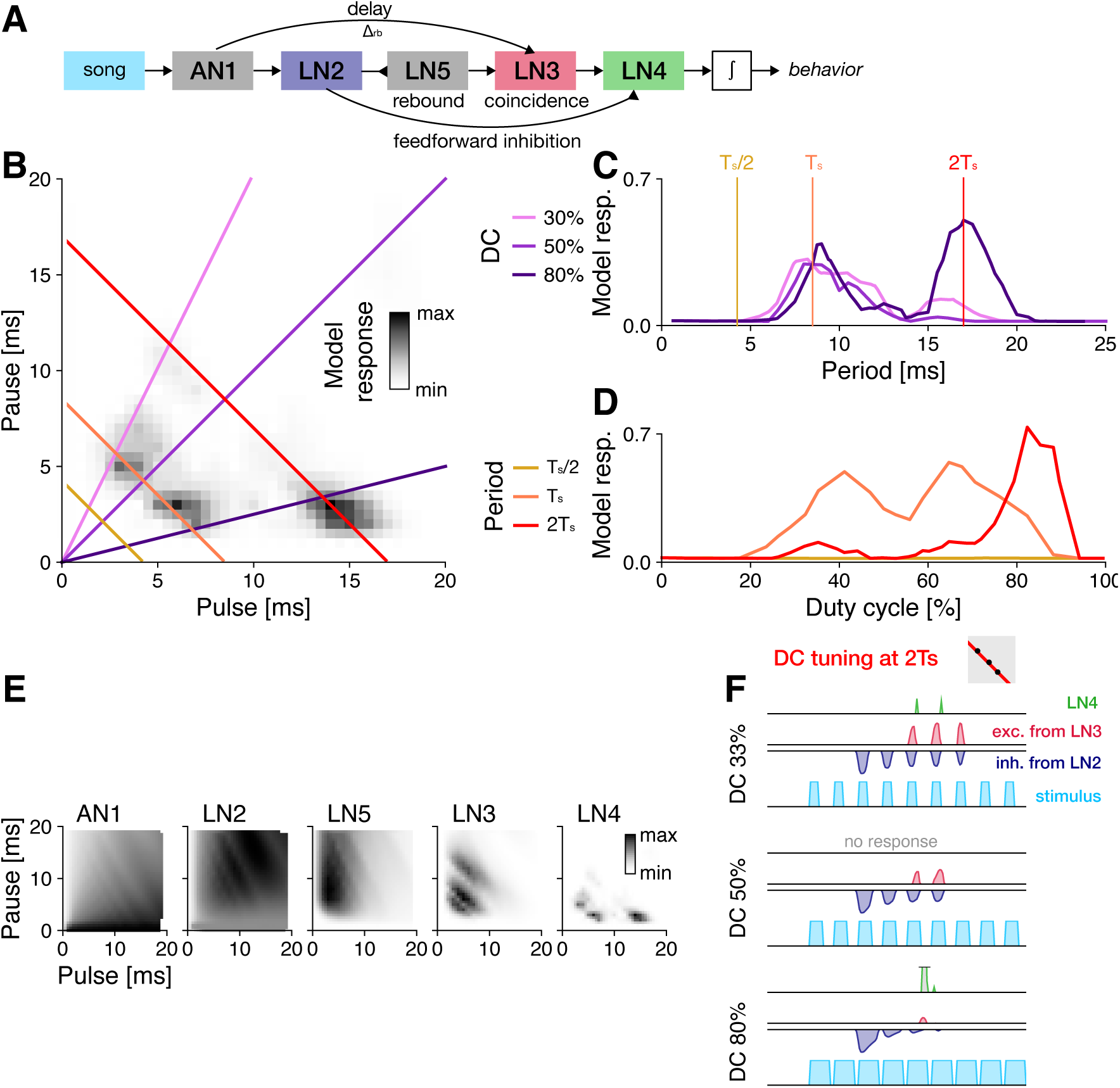
A model of the full song recognition network in crickets reproduces the resonant tuning of Anurogryllus. **A** Schematic of the full 5-neuron network and internal connections. Pointy and blunt ended arrows indicate excitation and inhibition, respectively. Delay (AN1-LN3), rebound (LN5), and coincidence (LN3) are computations of the core rebound mechanism (Fig. 3). Feed forward inhibition from LN2 to LN4 is crucial for reproducing DC tuning. **B** The resonant phenotype of Anurogryllus recovered with the five neuron model. Colored lines correspond the period and DC transects in D and E. **C** Traces of the DC transects labeled in B at 33%, 50%, and 80% DC, which each reveal the relative strength of the peaks at *T_s_* /*2*, *T_s_*, and *2 T_s_*. There is no response at the shortest period (*T_s_* /*2* —yellow). At the period of male song (*T_s_* —orange), DC tuning is band-pass. At the 17 ms period (*2 T_s_* —red), tuning is biphasic, as observed in the behavioral data. Vertical lines correspond to the DCs shown in C. **D** Traces of the period transects labeled in B, which shows that each peak has unique DC preferences (Compare with the behavioral data in Fig 1 which shows that Anurogryllus similarly demonstrates a bandpass preference around the male calling song *T_s_*, and a preference for high DCs for the *2 T_s_* peak). **E** Response profiles of the five neurons in the network. **F** Example internal network traces for three songs (blue) along the *2 T_s_* period transect at different DC’s, showing the interaction of the excitatory coincidence detection output from LN3 (red) and the inhibition from LN2 (blue) to produce the output response in LN4 (green). Gray boxes in F illustrate the stimulus parameters for which traces are shown in the context of the PPF (compare B).

### 2.3 The mother network can produce the resonant phenotype

A computational model of the song recognition network in crickets, that was originally constructed to reproduce electrophysiological data from *G. bimaculatus* (Clemens et al., 2021), was fitted to the behavioral data from Anurogryllus females (Fig. 1C). This model reproduced the Anurogryllus behavior (5A): Resonant peaks at *T_s_* and *2 T_s_*, and DC tuning at *2 T_s_* that is bimodal with a preference for higher DCs. This supports the mother network hypothesis—the network from *G. bimaculatus* can produce the preference profiles from all cricket species examined so far and could therefore constitute the template network for song recognition in crickets. How does the characteristic period and DC tuning arise in the network? Above, we have shown that the rebound mechanism at the core of the network is sufficient to produce resonant responses but insufficient to produce the DC tuning at *2 T_s_* (Fig. 3 D). We therefore investigated where in the network both response properties arise.

In the full network, LN3 is equivalent to the output of the simple rebound model as it is the coincidence detector that receives input from the rebound neuron LN5 and a delayed input from AN1. Accordingly, resonant tuning with responses at multiple periods in the network arises in LN3 (Fig. 5E). Indeed, the effective delay between the two inputs to LN3 is 25.3 ms, similar to the delay *Δ_rb_* = *23* ms found for the simple rebound model.

The DC tuning of Anurogryllus arises in the last neuron of the full network, in LN4 (Fig. 5E). LN4 receives excitatory input from the coincidence detector LN3 and feed-forward inhibition from LN2. The inhibition from LN2 shapes the DC tuning by suppressing responses to song with a DC of 50% at *2 T_s_* (Fig. 5F): At DCs around 50%, the excitatory input from LN3 is ineffectual because it overlaps with the strong inhibition from LN2. For higher and lower DCs, inhibition is less potent and hence the output from coincidence detection prevails and LN4 responds. At lower DCs, inhibition is weak and offset in time from the excitatory input from LN3. At higher DCs, the excitation from LN3 is stronger and arrives slightly later than the inhibition. In summary, the Anurogryllus tuning arises serially, through two computations in the network model: Rebound and coincidence detection in LN3 shape the period tuning and feed-forward inhibition from LN2 suppresses responses at wrong periods and shapes the DC tuning.

To confirm that a mechanism comprising rebounds and feed-forward inhibition is sufficient to reproduce the Anurogryllus behavior, we extended the simple rebound model (Fig. 3) with delayed feed-forward inhibition (Fig. S2A). We used the parameters of the simple rebound model (Fig. 3) and only fitted the delay and filter properties of the LN2-like input to LN4 (see Methods). This model is sufficient to reproduce the resonant period tuning (Fig. S2C) and the bimodal DC tuning of Anurogryllus (Fig. S2B–D). The DC tuning arises from the timing of excitatory and inhibitory inputs to LN4 (Fig. S2E), not from their strengths (Fig. S2F). Responses to a DC of 50% are suppressed in LN4 because excitation and inhibition arrive at the same time (Fig. S2E). For shorter/longer DCs, inhibition arrives too early/late to cancel the excitation.

## 3 Discussion

In this paper, we investigated the consequences of the unique song recognition phenotype of Anurogryllus for the evolution of acoustic communication among crickets. Anurogryllus females respond to three different pulse patterns: pulse patterns matching the period of the male song but also to patterns with twice the period and low or high duty cycle(Fig. 1). Using computational modeling, we tested whether this unusual recognition phenotype in crickets necessitates a corresponding novel recognition mechanism or if the hypothesized shared mechanism observed in other cricket species suffices.

First, to identify elemental computations required for resonant song recognition, we tested simple delay and filter-based network models, alongside a single-neuron model with resonant membrane properties (Figs. 2, 3, 4). While each model could resonate, it was the resonate and fire single-neuron model that best matched the tuning of Anurogryllus for both period and DC. That a single-neuron model qualitatively matches the behavior suggests that changes in intracellular properties capable of inducing oscillations of the membrane potential could underlie the evolution of resonant song recognition in Anurogryllus.

Critically, we found that a pure rebound mechanism, the core computation of the hypothesized shared mechanism for song recognition in crickets, is insufficient to reproduce the tuning of Anurogryllus and that additional computations present in the shared network are necessary (Fig. 5): The core rebound mechanism gives rise only to the resonant period tuning but not the DC tuning, casting doubt on the mother network hypothesis. However, the addition of feed-forward inhibition, present in the full model, recovered the DC tuning profile (Fig. S2). In *G. bimaculatus*, the cricket species in which the network was described, feed-forward inhibition primarily served to refine period tuning (Schöneich et al., 2015), as no resonances appear at the coincidence detection stage of the network in this case. In Anurogryllus, it appears to have been coopted to modulate DC tuning by attenuating responses to intermediate DCs. This is in agreement with the fitted mother network model, in which the resonant response arises in two steps: The rebound-based mechanism at the core of the network shapes the period tuning, while feed-forward inhibition shapes the DC tuning. Importantly, in the original network, the feed-forward inhibition only sharpens the period tuning, suggesting that this computation can be re-used in Anurogryllus for another function. Overall, our study shows how novel behaviors can arise from the modification of existing intracellular and network computations.

### Mechanisms of resonant song recognition in Anurogryllus

In the absence of physiological recordings, computational modeling can be used to constrain hypotheses about the recognition mechanism of Anurogryllus. Here, we used two approaches: 1) Minimal models of networks and single-neurons, to identify the computations required to produce the Anurogryllus tuning, and 2) a complex network model based on the song recognition network from another species, *G. bimaculatus*, to test about the potential of that network to produce resonant behavior. We identified two mechanisms that can give rise to the resonant song recognition in Anurogryllus: A cell-intrinsic mechanism based on oscillatory membrane properties (Fig. 4). And a combination of two network mechanisms: rebound and feed-forward inhibition (Fig. 5, S2).

The resonate and fire neuron, a single-neuron model that was previously used to reproduce resonant song recognition in a katydid (Webb et al., 2007) qualitatively reproduced the pulse rate and DC tuning of Anurogryllus. It produced responses to *T_s_* and *2 T_s_* and exhibited bi-modal DC tuning at *2 T_s_* (Fig. 4). Off-target responses in the model for longer periods could be suppressed by additional computations like a high-pass filter, for instance via adaptation, a computation that is ubiquitous in the mother network (Benda, 2021; Clemens et al., 2020). The resonant membrane properties could arise by changing the expression levels of specific ion channels in any of the neurons of the mother network. For instance, voltage-gated calcium (Ca_V_) or potassium (KCNQ, HCN) (Ge and Liu, 2016).

We also found that the rebound mechanism in the mother network alone was not sufficient to produce the tuning of Anurogryllus (Fig. 3). However, combining the rebound with feed-forward inhibition recreates the period and DC tuning (Fig. S2). In the full network model (Fig. 5A, B), these computations arise in different neurons of the network: First, the rebound produced by LN5 is combined with delayed excitation from AN1 in the coincidence detector LN3 to produce the resonant period tuning. Then, feed-forward inhibition from LN2 shapes the DC tuning in LN4. Crucial for tuning the network are the response delays: from AN1 onto LN3 to tune the preferred periods (Clemens et al., 2021) and from LN2 onto LN4 to tune the DC responses (Fig. S2E, F). Ultimately, determining which of the two proposed mechanisms—single cell or network—generates the resonant behavior of Anurogryllus will require intracellular recordings that detect membrane oscillations and assess response delays.

### Resonances are rare because they are undesirable, or because they have been missed in experiments

Our analysis of the simple models revealed that resonances can arise easily from common mechanisms like delays or membrane oscillations (Fig. 2, 3, 4). However, multi-peaked response profiles are known only from two species—Anurogryllus and *T. cantans* (Bush and Schul, 2006). This may reflect selection against resonant tuning, because resonances broaden the female tuning to regions that fall outside of the male calling song, leading to the potential misidentification of mating partners. While multi-peaked tuning can still enable mate recognition if other signalers do not sing at the resonant off-target peaks (Amézquita et al., 2011), it is likely that these resonances are suppressed and hidden in many pattern recognition networks.

However, multi-peaked responses might also be underreported, since their detection requires a comprehensive and systematic sampling of the stimulus space when quantifying female preferences. Future playback experiments should therefore be designed to ensure the detection of resonances: Stimuli should not only densely sample different periods but should also do so at multiple DCs. For instance, a stimulus set that densely samples pulse periods, but at a DC of 50% would have missed the resonant peaks at twice the song period in Anurogryllus and *T. cantans* (Fig. 1G, H). A characterization of the DC tuning also helps differentiate between resonant mechanisms (Figs. 2, 3, 4): Only the R&F model produces bimodal DC tuning at *2 T_s_*, while autocorrelation and rebound mechanisms produce unimodal DC tuning. Sweep or chirp stimuli commonly used in electrophysiology have a changing pulse rate or period are not sufficient for discriminating models since these stimuli have a constant DC (Narayanan and Johnston, 2007). Similarly, the presence or absence of responses to odd and even multiples or fractions of the song period can disambiguate between different mechanisms (Figs. 2, 3, 4): The R&F model responds only to even multiples of the model’s characteristic period, while the simple delay-based models respond to both even and odd fractions. However, an interpretation of such experiments is complicated by the fact that the behavioral preference is the outcome of multiple computations, in the case of Anurogryllus possibly of a rebound mechanism combined with feed-forward inhibition (Fig. S2).

### Nonlinear computations accelerate the evolutionary divergence of song preferences

While having potentially negative consequences for species recognition, neuronal resonances could support the fast divergence of behaviors: According to the standard model of evolution, novel phenotypes evolve through an accumulation of small behavioral changes. This gradual change is opposed to saltatory changes, which poses that nonlinearities in the mapping from genotype (or system parameter) to phenotype can drive sudden large phenotypic changes (Gould and Eldredge, 1977). Evolutionary developmental biology has shown that strongly nonlinear developmental programs can give rise to morphological innovations from small genetic changes (Müller, 2007)—so-called morphological monsters. Resonant song recognition with responses to disjoint sets of songs is also the result of a highly nonlinear mapping from network parameters to behavior. If simple mechanisms existed to isolate individual peaks, then behavioral preferences could jump between these resonant peaks. The resulting step-like changes in female preference will in turn drive strong changes in male song and a rapid isolation between sister species. Spike-frequency adaptation (SFA) is ubiquitous in the nervous system and is also found in the song recognition network of *G. bimaculatus* (Clemens et al., 2021). SFA in combination with the low-pass properties of the membrane results in a band-pass filter that can be tuned by changing the time constants of the membrane or of the adaptation current (Benda and Herz, 2003; Benda, 2021). We have implemented a simple proof-of-principle model to illustrates that SFA can isolate individual peaks from a resonant response field (Fig. SS3). Changes in parameters will change the relative amplitudes of the individual peaks without creating intermediate peaks. This will exert selection pressure on the male song to jump to the new larger peak of the female preference function. Thus, strong nonlinearities in the mapping from genotype to behavior—like resonances—can drive rapid evolutionary change.

## Conclusion

Overall, our computational approach revealed the capacity of neural networks for change: The song recognition network described in *G. bimaculatus* consists of a set of elementary computations— rebounds, coincidence detection, adaptation, feed-forward inhibition—that can give rise to a rich set of recognition behaviors. This network has the capacity to produce all recognition types known in crickets: For pulse pause, pulse period, DC, and even for the “behavioral monster” Anurogryllus, with its complex resonant tuning. This network could therefore serve as a mother network, that gives rise to the full diversity of song recognition in crickets. That even a small network, consisting of only 5 neurons can produce so many diverse behaviors highlights the enormous potential of neural networks to produce evolutionary novel phenotypes.

While simple models of the core rebound, delay, and coincidence detection mechanism only partially recovered the characteristics of the resonant Anurogryllus behavior, insights from the full model revealed that the function of the pulse recognition network in crickets might include additional selectivity for duty cycle via feed-forward inhibition, the inclusion of which enabled the simple rebound model to replicate the resonant pattern. These results further suggest that the function of the pulse pattern recognition network in crickets cannot be conceptualized merely as a rate detector, but that it may additionally select for the duty cycle characteristics of incoming song, necessitating the inclusion of feed-forward inhibition in even a minimal model for song feature recognition networks in crickets. More generally, the capacity of neuronal networks to drive evolutionary change stems in part from the multitude of nonlinear computations at the network and single-neuron level, which can be coopted to produce new behaviors.

## Acknowledgments

We thank the following students who ran the behavioral experiments for establishing the Anurogryllus female response: Sofia Hayden, Daria Ivanova, Eileen Gabel, Kolja Haß, Anne Görlitz. Funded by the Deutsche Forschungsgemeinschaft (DFG, German Research Foundation) as part of the SPP 2205, project number 430158535.

## Contributions

- Conceptualization – WM, MH, JC
- Animals and behavioral experiments – BE, MH
- Modeling and analysis – WM, JC
- First draft – WM, JC
- Feedback on draft – BE, MH

## 4 Methods

### 4.1 Animals

Behavioral experiments were performed with *Anurogryllus muticus* from the same colony as used in (Erregger et al., 2018). The progeny of individuals caught on Barro Colorado Island in Panama were reared to adulthood at the Department of Zoology at the University of Graz in Austria and held at 25–28 °C with *ad libitum* food and water. Starting with the second or third instar, individuals were separated from the colony and placed in individual plastic boxes.

### 4.2 Male song recording and analysis

Individuals were placed in an array of separate boxes (mean temperature 24.9*±*1.0°C SD) for a duration of 16–24 hours. Each box was equipped with a microphone and isolating foam to ensure acoustic isolation. Using customized software (LabVIEW 7, National Instruments, Austin, TX, USA), the microphone (TCM 141 Conrad; Conrad Electronic, Germany) in each box was scanned for 800 ms at a time with a sampling rate of 100 kHz and a male was recorded for 20 s if it produced sound during that 800-ms interval (Hennig et al., 2016). The song carrier frequency was determined from the spectral peak of the raw waveform signal. For analysis of the temporal pattern, the normalized envelope of the song signal was computed after signal rectification by squaring and low-pass filtering at 200 Hz (equivalent to a temporal resolution of 2.5 ms). Temporal parameters such as pulse and pause duration were calculated when the envelope crossed or fell below a threshold value at 10–15% of the signal envelope.

### 4.3 Female preference functions

Female preference was tested using a trackball system as described in (Hennig et al., 2016). Females, mounted to a metal rod, were placed on a hollow Styrofoam sphere (diameter: 100 mm, weight: 1.2–1.8 g) supported by an air stream between two perpendicularly placed loudspeakers (Piezo Horntweeter PH8; Conrad Electronic) in a wooden box with sound absorbing foam. Each loudspeaker was calibrated with a Bruel and Kjaer 2231 sound level meter and a half-inch condenser microphone (Bruel and Kjaer 4133 relative to 0.02 mPa, fast reading) at the top of the sphere where the female cricket was placed during experiments.

Digitally stored sound signals were transmitted from a hard disk by a D/A-board (update rate: 100 kHz, PCI 6221; National Instruments, Austin, TX, USA) to a digitally controlled attenuator (PA5; Tucker-Davis, Alachua, FL, USA), amplified (Raveland; Conrad Electronic) and broadcasted through the speakers. The longitudinal and lateral movements of the sphere were recorded by either a single optical sensor (Agilent ADNS-2051; Agilent Technologies, Santa Clara, CA, USA) at the bottom of the half-sphere or by two sensors (ADNS-5050; Avago Technologies, San Jose, CA, USA) with a focusing lens positioned laterally at an angle of 90°.

A silent control was used to monitor baseline walking activity, and a continuous tone was used to control for motivation and selectivity of female responses. At the beginning and the end of each test session, a species-specific, attractive song signal was presented to control for possible changes in phonotactic motivation during a session. For each test signal, the lateral deviation of a female during signal presentation for each of the two speakers was averaged and normalized with respect to the attractive control signal. The resulting phonotactic scores were therefore typically between 0 (no orientation towards the sound signal) and 1 (strong orientation towards the signal), although negative scores (orientation away from the signal) and scores higher than 1 (orientation towards signal stronger compared to control stimulus) were possible. Test signals and controls were presented at 80 dB sound pressure level. All tests included the four control stimuli (silent, continuous tone, and an attractive stimulus at the beginning and end of a test) and eight test stimuli (total duration was 29 min per test), and were performed at 24°C.

Phonotaxis values were measured for 75 artificial pulse trains, split into 10 playlists. Each playlist was tested with 3-8 females and the phonotaxis values for each stimulus were averaged over the females (Fig. S1. All stimulus parameters, phonotaxis values, and number of animals are listed in a supplemental data file. From the 75 average phonotaxis values, we generated a two-dimensional preference function using natural neighbor interpolation implemented in metpy (URL: https://github.com/Unidata/MetPy). The preference function covered pulse and pause durations between 0 and 20 ms, with a resolution of 0.1 ms. Negative phonotaxis values in this interpolated preference functions were set to 0.

### 4.4 Modeling

#### Stimulus and response data

Song pulses were constructed as rectangular boxes with an amplitude of 1. While natural pulse trains in Anurogryllus last for many seconds, the models tested here have dynamics on the timescale of a few tens of milliseconds. To speed up simulations, we therefore used pulse trains with a duration of 400 ms and omitted onset and offset transients when translating the model output to predicted phonotaxis (see below). The stimulus set contained pulse trains with all combinations of pulse and pause durations between 0–20 ms sampled on a grid with an interval of 0.5 ms, totalling (*20* /*0.5*)*^2^* = *1600* stimuli. As the fitting target, we used the two-dimensional preference function from Anurogryllus females obtained by interpolating the experimental phonotaxis values as described above, but on a grid with a step size of 0.5 ms.

#### Predicting phonotaxis score from model responses

The predicted phonotaxis score, *p*, is given by the average model response *r* (*t*) over the stimulus duration *D_s_*, excluding the first 25 ms and the last 10 ms to reduce the impact of response transients: 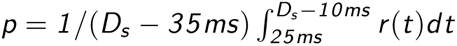.

#### Model fitting

The models were fitted using the Nelder-Mead method implemented in scipy.optimize.minimize, by minimizing the mean-squared error between the interpolated phonotaxis values from the data and the model response. If not stated otherwise, initial values for all parameters were set using a vector of initial conditions chosen manually to speed up fitting. Fits were run multiple times from slightly different initial conditions to avoid getting stuck in local minima. The presented parameter values are from models with the lowest error. The model parameters for the simple models are listed in Table 2 and for the full network in Table 3. The code and parameters for running all models can be found at https://github.com/janclemenslab/anurogryllus-monster.

**Table 2:**
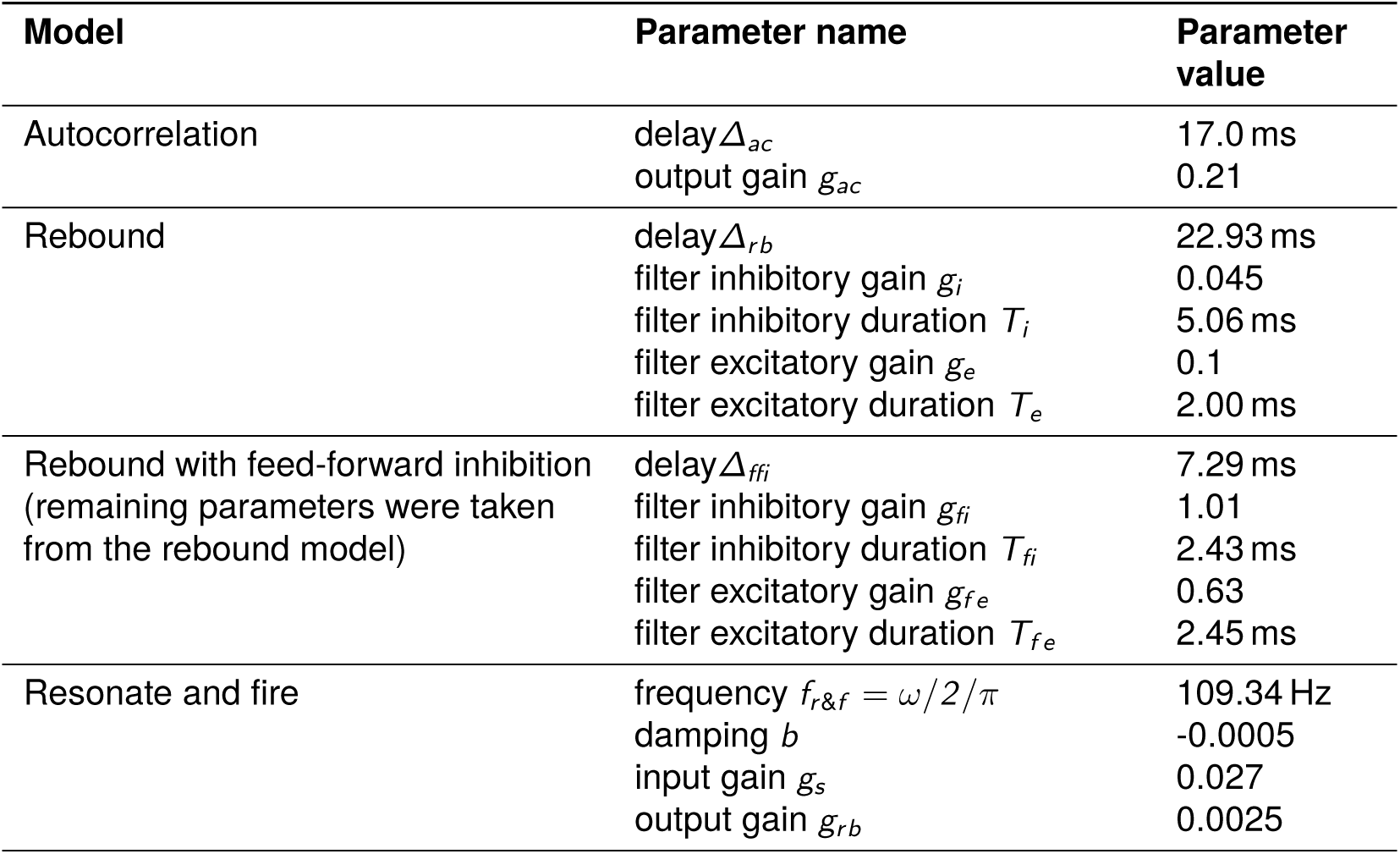
Parameters of the simple models fitted to reproduce the Anurogryllus preference function.

**Table 3:**
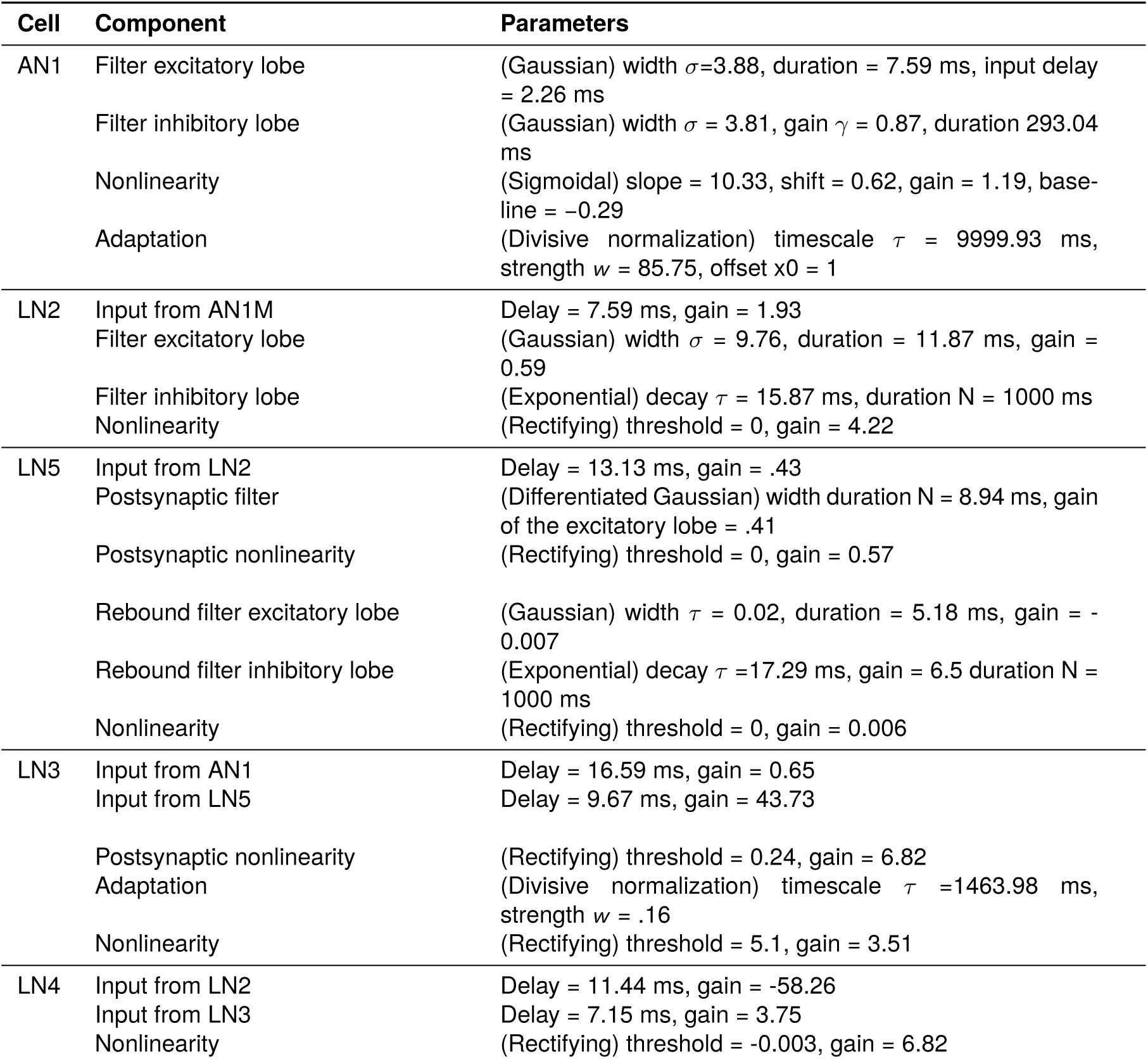
Parameters of the 5 neuron “mother network” model fitted to reproduce the Anurogryllus preference function.

**Table 4:**
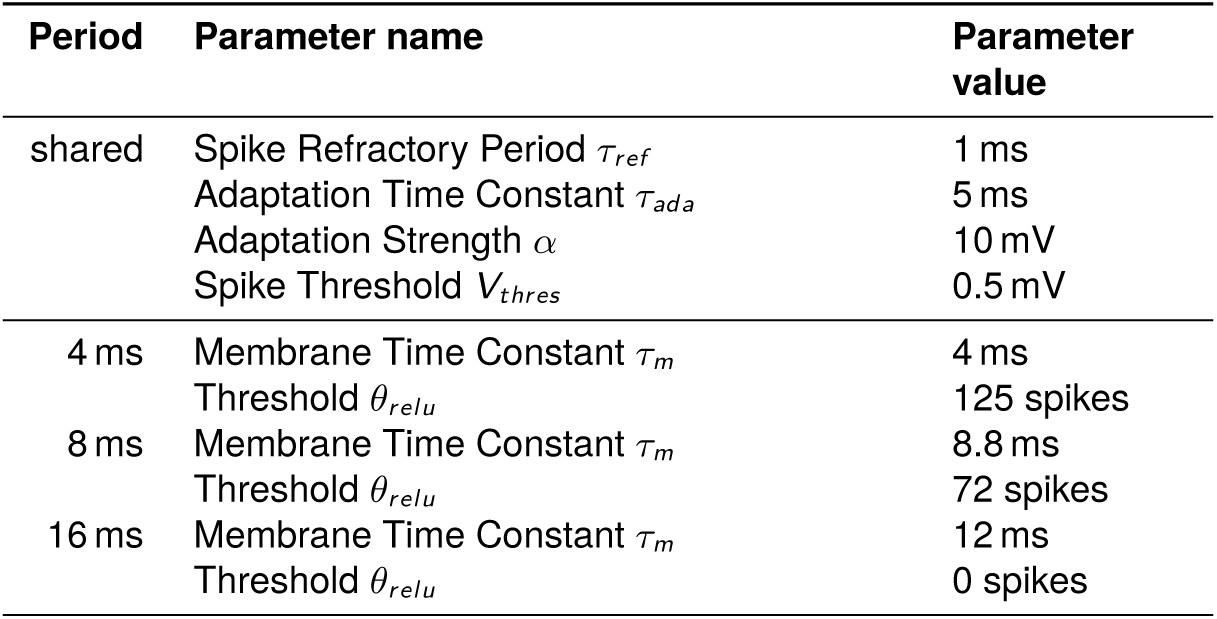
Parameters of the rebound model with adaptation shown in Fig. S3. The first four parameters in the table are shared between all variants of the model. Resonant peaks in the input from the rebound model at 4, 8, and 16 ms are isolated by adjusting the membrane time constant and the threshold of the rectifying linear function applied to the integrated output of the LIFAC model neuron.

**Table 5:**
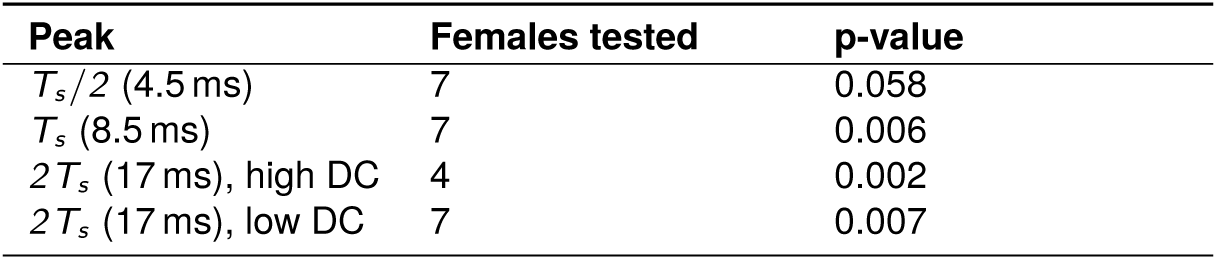
Statistical tests for each peak in the Anurogryllus phenotype. P-values were obtained from a paired one-sided t-test testing the hypothesis that the responses of the individuals to songs at the peak are greater than a silent control. All peaks, except for the peak at *T_s_* /*2*, are significant. The broad peak at *2 T_s_* for low DC was evaluated using two points within this peak for which different sets of females were tested. The stimuli for these low DC points have either a pause of 12.5 ms and a duration of 4.5 ms, or a pause of 11.2 ms and a duration of 5.8 ms. The high DC condition for *2 T_s_* was evaluated at a pause of 2.8 ms and a duration of 14.2 ms.

#### Autocorrelation model

In the autocorrelation model (Fig. 2A), the stimulus *s*(*t*) is delayed by *Δ_ac_*, *s_Δ_*(*t*) = *s*(*t − Δ_ac_*). A coincidence detector then multiplies *s*(*t*) and *s_Δ_*(*t*) and scales the result with a gain factor *g_ac_*: *r* (*t*) = *g_ac_ · s*(*t*) *· s_Δ_*(*t*). We did not add a nonlinearity to the output *r* (*t*), like a sigmoidal, prior or after integration, since it did not produce quantitatively different predictions during fitting. The autocorrelation model was simulated with a resolution of 10 kHz.

#### Rebound model

The rebound model extends the autocorrelation model by inverting and filtering one of the two paths the stimulus takes before coincidence detection to produce offset responses at the end of each pulse: 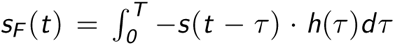. The filter 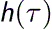 consists of two lobes, defined as rectangular windows: An inhibitory lobe with negative gain *g_i_* and duration *T_i_*, followed by an excitatory lobe with positive gain *g_e_* and duration *T_e_*. The positive response components in *s_F_* (*t*) corresponding to the rebound are isolated using a rectifying linear function: *s_R_* (*t*) = *f* (*s_F_* (*t*)), where *f* (*x*) = *0* if *x ≤ 0*, and *f* (*x*) = *x* if *x* = *0*. The coincidence detector then multiplies *s_R_* (*t*) and *s_Δ_*(*t*): *r* (*t*) = *s_R_* (*t*) *· s_Δ_*(*t*). The rebound model was simulated with a resolution of 4 kHz to accelerate the fitting process.

#### Rebound model with feed-forward inhibition

The rebound model with feed-forward inhibition extends the simple rebound model by including an additional inhibitory connection to the basic rebound model following coincidence detection (See Fig S2A). The added inhibitory path from stimulus to output (LN4) contains a bi-phasic filter with rectangular negative and positive lobes (similar to the filter in the rebound model) and a delay. The negative components of the output of the bi-phasic filter were then used as an inhibitory input to an LN4-like output neuron. The LN4-like neuron combines the inputs from the coincidence detector and the feed-forward inhibitory paths. To obtain the predicted phonotaxis value for a given stimulus, the output of the LN4-like neuron was passed through a rectifying linear function with threshold *θ_relu_* = *0* and a linear gain *g_relu_* = *1* and then integrated. When fitting this model, the parameters of the simple rebound model fitted previously were kept fixed and only the additional parameters for the feed-forward inhibition branch (the delay time and the gain and duration of the inhibitory and excitatory lobe) were adjusted.

#### Resonate-and-fire neuron

The resonate-and-fire model was implemented following Izhikevich (2001):

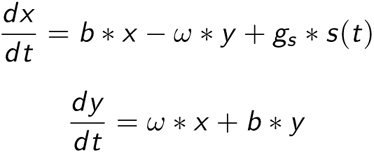

where *x* is a current-like variable, *y* is a voltage-like variable, *b* is the damping factor, *ω* is the intrinsic frequency, *s*(*t*) is the song input and *g_s_* is the gain of the song input. If *y* exceeds the threshold *y_threshold_* = *1*, a spike with amplitude *g_rb_* is elicited and current and voltage are reset to *x_reset_* = *0* and *y_reset_* = *1*:

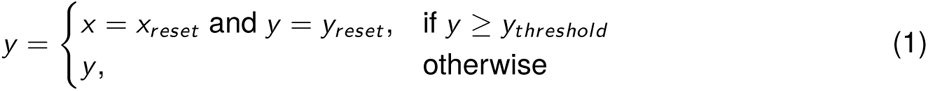

The differential equations were numerically integrated using the Euler method with a time step of 0.1 ms.

Full model of the song recognition network in *G. bimaculatus*

To test whether the song recognition network from *G. bimaculatus* described in Schöneich et al. (2015) can reproduce the resonant behavior of Anurogryllus, we used the model of the network from Clemens et al. (2020). This model was fitted to reproduce the response dynamics and the tuning of all neurons in the network using electrophysiological recordings from*G. bimaculatus* for a large set of pulse train stimuli (Kostarakos and Hedwig, 2012; Schöneich et al., 2015). The forty-five parameters in the network model were fitted using the Nelder-Mead optimization algorithm, by minimizing the mean-square error between experimental and predicted phototaxis values (see Table 3 for the fitted parameters) using the parameter values found for *G. bimaculatus* as a start point. Several rounds of optimization were required to converge on the given parameter set, with Gaussian-distributed noise added to all parameters at the start of the initial optimization rounds to avoid undesirable local minima. Model fitting often yielded models that reproduced the tuning of Anurogryllus with only transient responses at the onset of the pulse train. Given that Anurogryllus song lasts multiple seconds and elicits phonotaxis throughout, we deemed these solutions physiologically unrealistic. We therefore added the constraint that responses of AN1 in the model should spike throughout the stimulus for pulse trains with conspecific parameters.

#### Modeling jumps between resonant peaks with spike-frequency adaptation

To demonstrate that individual resonant peaks can be isolated from a resonant response field, we added to the rebound model fitted to the Anurogryllus data (Fig. 3, same parameters as in Table 2) a leaky integrate and fire neuron with an adaptation current (LIFAC) using the code published with Benda (2021). The LIFAC model is driven by the non-integrated output of the rebound model and acts as band-pass filter, because it combines the low-pass properties of a cell membrane and high-pass properties from adaptation (Benda and Herz, 2003). The total spike output from the LIFAC model for each stimulus is passed through a rectifying linear function with threshold *θ_relu_* and a linear gain *g_relu_* = *1*, to compute the predicted phonotaxis value.

The LIFAC neuron responds to a current input *I* by increasing the membrane potential *V* from which an adaptation current *A* is subtracted:

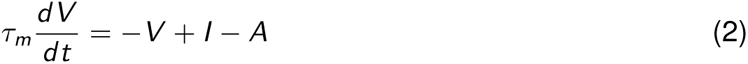

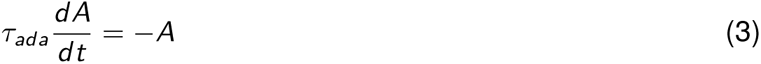

with time constants of the membrane and of adaptation, *τ_m_* and *τ_ada_*, respectively. If the voltage *V* reaches the spiking threshold *V_thres_*, a spike is elicited, and *V* is reset to *V_reset_* and the adaptation current strength *A* is incremented by *α*:

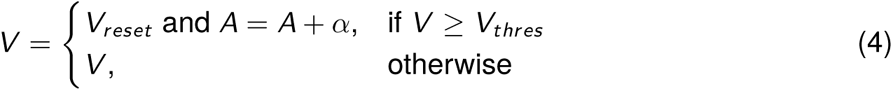

Each spike initiates a refractory period *τ_ref_*, during which both *V* and *A* are fixed to their reset values.

## 5 Supplemental material

**Figure S1:**
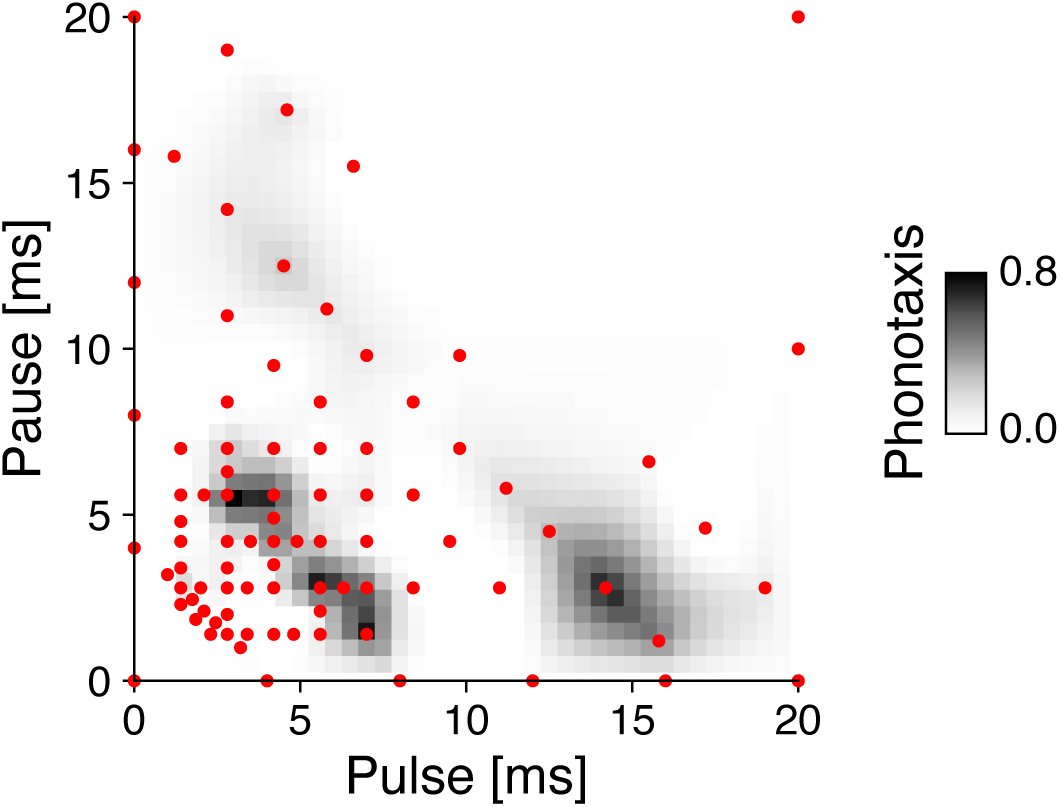
Pulse train stimuli used for estimating the pulse-pause field (PPF) shown in Figure 1C. Individual pulse trains for which phonotaxis values were measured are shown as red dots. The PPF (color coded, see color bar) was obtained by natural neighbor interpolation of the phonotaxis values on a dense 41×41 grid (all combinations of pulses and pauses between 0 and 20 ms with a step size of 0.1 ms). Phonotaxis values at the boundaries (pulse or pause 0 ms) were set to 0.

**Figure S2:**
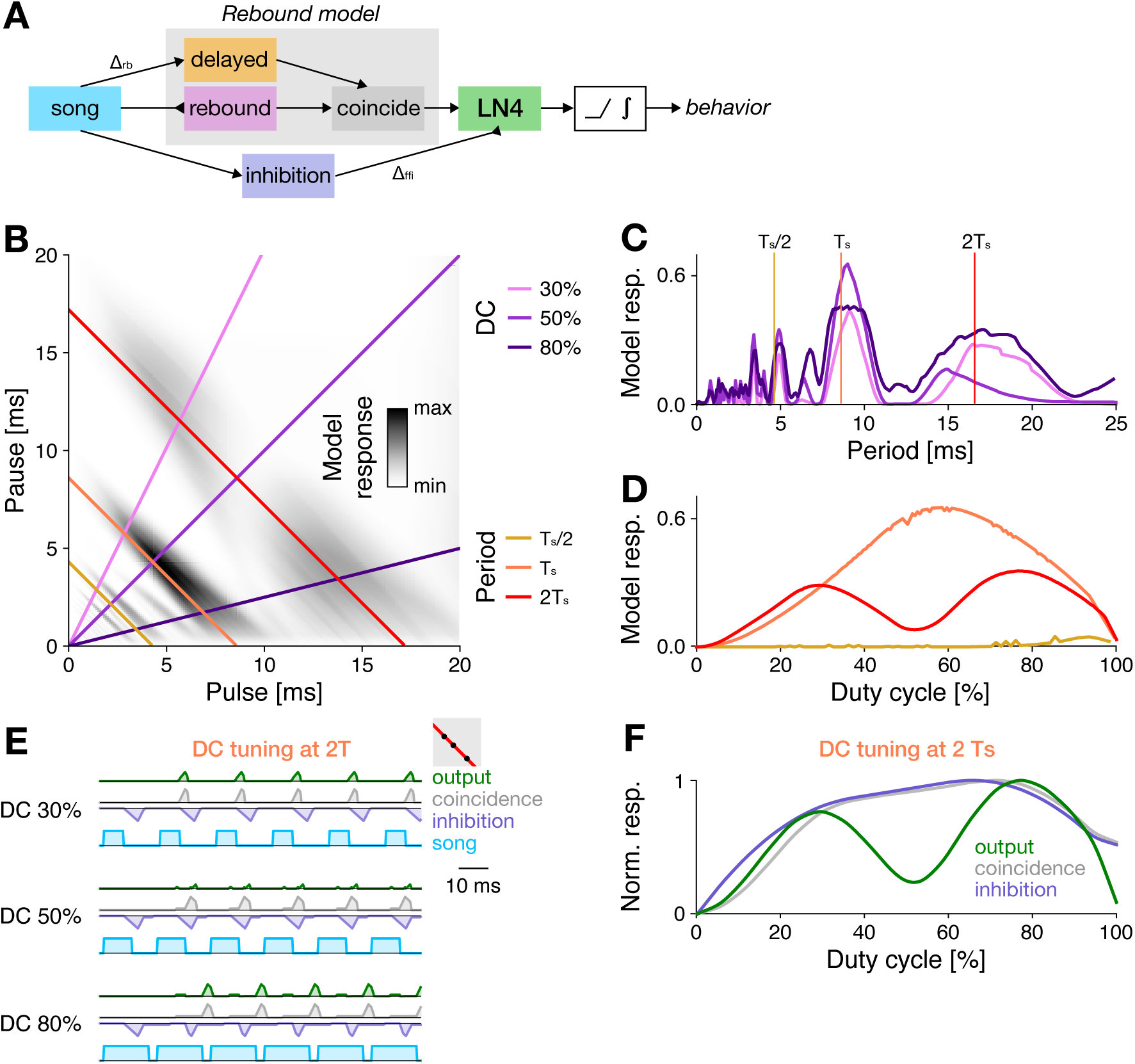
A combination of rebound and feed-forward inhibition are sufficient to create the tuning of Anurogryllus. **A** Schematic of the rebound model with delayed feed-forward inhibition. An LN4-like neuron receives input from the coincidence detector of a rebound model and from inhibition. The output of **B** PPF illustrating the responses produced by the modified rebound model fitted to behavioral data from Anurogryllus (Fig.1C, see color bar), demonstrating the restored bimodal shape of the 17 ms period transect. Colored lines correspond to the DC and period transects shown in C and D. **C** Period tuning of the model for different DCs. **D** DC tuning for three different pulse periods, corresponding to *T_s_* /*2*, *T_s_*, and *2 T_s_*. The curves indicate bandpass preference around the male calling song *T_s_*, and bimodal DC tuning for the *2 T_s_* peak. Vertical lines correspond to the DCs shown in C. **E** Example traces showing how the delay timing of inhibition (blue) interacts with the coincidence detection output (grey) to produce bimodal tuning along the *2 T_s_* 17 ms period transect. Inhibition at 50% DC coincides with the timing of the coincidence detection output, fully suppressing responses. **F** DC tuning of the rebound output (grey) vs the feed-forward inhibition (blue) for the 2T 17 ms transect, which produces the final bimodal tuning (green) as observed in the behavior data.

**Figure S3:**
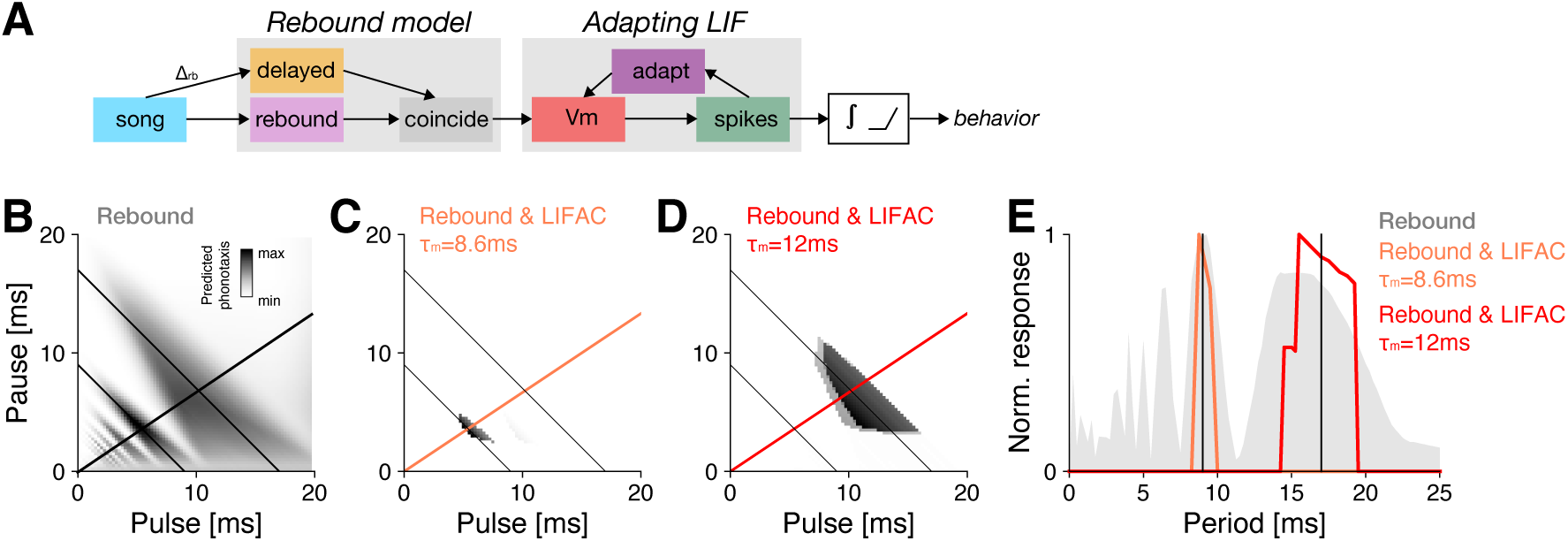
Hidden resonances enable saltatory evolution of song preferences. **A** Structure of the rebound model with adaptation. The non-integrated output of the rebound model from Fig. 3 (see B) was used to drive a leaky integrate and fire neuron with an adaptation current (LIFAC). The spike output of the LIFAC is then integrated to yield a value proportional to the phonotaxis. A rectifying nonlinearity (relu) is then used to further sharpen the tuning for song. The LIFAC acts as a band-pass filter that was tuned via the time constant of the membrane *τ_m_*. **B** PPF of the rebound model with resonant peaks used as the input to the LIFAC. The two resonant peaks at *≈ 9* ms and *≈ 17* ms that were isolated with adaptation are shown as thin black antidiagonal lines. The thicker black diagonal line shows the transect at a DC of 66% shown in E. **C, D** PPFs of the rebound & LIFAC model. The membrane time constants *τ_m_* were tuned to 8.6 (C) and 12.2 ms (D) to isolate the resonant peaks at *≈ 9* ms and *≈ 17* ms (thin black lines). The thicker orange and red diagonal lines show the transects at a DC of 66% shown in E. **E** Period tuning of the models in B–D for a transect through the PPF at a DC of 66%.

